# Critical point drying of brain tissue for X-ray phase contrast imaging

**DOI:** 10.1101/2025.09.02.673701

**Authors:** S. Khan, J. Albers, A. Vorobyev, Y. Zhang, J. Reichmann, A. Svetlove, F. De Marco, K. Denisova, Y. Yang, F. Seichepine, J. O. Douglas, E. Duke, P. Cloetens, A. Pacureanu, A. T. Schaefer, C. Bosch

## Abstract

X-ray phase contrast tomography is emerging as a powerful method for imaging large volumes of brain tissue at sub-cellular resolution. However, current sample preparation methods are largely inherited from visible light or electron microscopy workflows and hence are not optimised to exploit the full potential of X-ray contrast mechanisms. Here we propose to replace interstitial material by air to enhance X-ray phase contrast of the ultrastructural features. We used critical point drying (CPD) of heavy metal-stained mouse brain tissue to produce mechanically stable samples with preserved ultrastructure and enhanced refractive index boundaries, a nanofoam-like material that remains compatible with follow-up conventional resin embedding. Using two complementary synchrotron-based setups, a high-throughput microtomography beamline (P14, DESY) and a nanoscale holographic tomography beamline (ID16A, ESRF), we found that CPD samples consistently showed 2–4*×* stronger phase-shift signal than conventional resin-embedded tissue. The contrast gain remained consistent across samples, imaging conditions, and beamlines. Our results suggest that CPD offers a versatile route for preparing tissue for subcellular and ultrastructural-resolution X-ray imaging. It retains structural detail while improving signal, and is compatible with other processing procedures like femtosecond laser milling or electron microscopy, paving the path for biological tissue imaging beyond the mm^3^ scale.

## Introduction

X-ray phase contrast imaging can efficiently map millimiter-scale volumes of soft tissue, a size that can contain complete modules of neuronal circuits in the mammalian brain, at subcellular resolution without the need to slice the samples, by leveraging the penetrating power, contrast mechanisms and flux of synchrotron-based coherent X-ray microscopy [1–4]. X-ray phase contrast imaging [5–8] offers the potential to bridge scales, capturing cellular and subcellular architecture, across intact tissue volumes with isotropic resolution approaching sub-40 nm detail [3, 9]. However, sample preparation protocols of biological tissues for X-ray phase contrast imaging [2, 4, 10–12] have largely been inherited from visible light and volume electron microscopy (vEM) workflows, including the resin embedding protocols following heavy metal staining. While these methods provide mechanical stability and compatibility with EM, they were not tailored to enhance X-ray contrast.

In the hard X-ray regime (5 - 100 keV) [13] which is typically used for imaging metal-stained biological tissues, image contrast arises from differences in refractive index between components. Upon interaction of an X-ray wavefront with matter, phase shifts distort the wavefront. Self-interference between refracted and un-refracted portions of the wavefront results in interference patterns that are measured as intensity variations after free-space propagation. Differently scattering neighbouring tissue regions thereby trigger different phase shifts to the wavefront that generate interference patterns (or fringes) at the interface of those regions, which ultimately give rise to contrast in phase-sensitive imaging [6, 7, 14]. In conventionally embedded tissue, contrast is generated by differences in phase shift and attenuation between heavy metal-stained structures and the surrounding carbon-rich embedding resin. Since the contrast stems from changes in electron density, one should expect to obtain enhanced contrast when replacing the interstitial material with vacuum /air instead of infiltrating it with an epoxy resin.

We therefore explored an alternative sample preparation route that could allow avoiding the embedding material altogether. Following heavy metal staining of the mouse brain tissue samples and dehydration with a series of incubations in increasing ethanol concentrations, we used critical point drying (CPD) to remove the ethanol from the samples. This yielded a mechanically stable nanofoam-like dry scaffold in which the metal-stained ultrastructure in the tissue was surrounded by air or vacuum, depending on the imaging requirements [15–18].

In CPD samples phase contrast should be constrained by the refractive index of the foreground, similar as in the conventional workflow (heavy metal-stained lipids and proteins present in plasma membranes and in protein clusters), and the refractive index of the background, which in this case should be largely minimised or negligible. This should result in more phase contrast generated by ultrastructural features in CPD-prepared samples than in resin-embedded counterparts.

We first validated that the ultrastructure of CPD-treated tissue remains preserved by embedding dried samples (post-CPD) in epoxy resin and examining them with transmission electron microscopy (TEM). We then carried out X-ray phase contrast imaging on matched sets of CPD and resin-embedded samples at two synchrotron instruments: the high-throughput imaging setup at the P14 beamline at DESY (PETRA III), and the nanoscale holographic tomography ID16A beamline at ESRF.

## Results & Discussion

For X-ray imaging of biological tissues to be useful for mapping ultrastructural details and long-range cellular tracing (such as for connectomics applications), it is essential that tissue ultrastructure remains well preserved through all stages of sample preparation. Any enhancement in phase contrast or imaging resolution is only valuable if the underlying biological features, such as cell boundaries (cytoplasmic membranes), synapses, and organelles, retain their morphology. To address this, we assessed whether CPD compromises ultrastructural integrity in a way that could undermine its broader applicability. We embedded CPD-treated samples in resin, sectioned them with standard ultramicrotomy tools, and imaged them using transmission electron microscopy (TEM).

Both methods to prepare samples (Fig. 1c, d) start with chemical fixation and heavy metal staining [4] of tissue slices obtained from the olfactory bulb of the mouse brain (see Methods). These initial steps follow established protocols for brain tissue preparation for volume electron microscopy and more recently X-ray based imaging for connectomics [1–4]. For standard resin embedding, after dehydration through graded ethanol series, samples are embedded in Epon (Fig. 1c,e). In the CPD workflow, ethanol-dehydrated samples were transferred into liquid CO_2_ and processed through the critical point (Fig. 1d,f-h). CPD replaces ethanol with liquid CO_2_ (Fig. 1g), which is brought to its critical point to enable drying with minimal surface tension effects, thereby reducing the risk of damage to fragile ultrastructural features [15–18]. The result is a dry, unsupported tissue sample with preserved structural integrity but with the mechanical properties of a metallic nanofoam [20] (Fig. 1f,i, Supplementary Fig. S1). Finally, after completing the respective preparation workflows for resin-embedded and CPD-treated tissue, cylindrical pillars of varying diameters were fabricated using femtosecond laser milling[21] (Supplementary Fig. S2, Tables 1 and 2).

**Table 1.** Summary of samples imaged using X-ray phase contrast tomography (XPCT) at beamline P14 (PETRA III, DESY). Samples were stained with heavy metals (rOTO protocol) and prepared using either critical point drying (CPD) or resin embedding (Epon). Imaging parameters include X-ray energy, effective pixel size, and exposure time.

| | sampleID | Paper Name | Staining | Embedding | Pillar diameter ($\mu\text{m}$) | Scan-Parameters | Figures |
| --- | --- | --- | --- | --- | --- | --- | --- |
| 1 | SK101_p1 | Epon1 | rOTO | Epon | 500 | E=17 keV; pixel size=325 nm; exposure time= 10 ms | Fig.3 |
| 2 | SK101_p3 | Epon2 | rOTO | Epon | 500 | E=17 keV; pixel size=325 nm; exposure time= 10 ms | Supp. Fig. S4 |
| 3 | SK106_p1 | Epon3 | rOTO | Epon | 500 | E=17 keV; pixel size=325 nm; exposure time= 10 ms | Supp. Fig. S4 |
| 4 | SK277_p1 | CPD1 | rOTO | CPD | 500 | E=17 keV; pixel size=325 nm; exposure time= 10 ms | Fig.3 |
| 5 | SK271_p1 | CPD2 | rOTO | CPD | 500 | E=17 keV; pixel size=325 nm; exposure time= 10 ms | Supp. Fig. S5 |
| 6 | SK282_p1 | CPD3 | rOTO | CPD | 500 | E=17 keV; pixel size=325 nm; exposure time= 10 ms | Supp. Fig. S5 |
| 7 | SK287_p1 | CPD4 | rOTO | CPD | 500 | E=17 keV; pixel size=325 nm; exposure time= 10 ms | Supp. Fig. S5 |
| 8 | SK197_p1 | Paraffin | Unstained | Paraffin | 500 | E=17 keV; pixel size=325 nm; exposure time= 10 ms | Supp. Fig. S7 |
| 9 | SK207_p1 | CPD | Unstained | CPD | 500 | E=17 keV; pixel size=325 nm; exposure time= 10 ms | Supp. Fig. S7 |

**Table 2.** Summary of samples imaged by X-ray nano-holotomography (XNH) at beamline ID16A (ESRF). Each sample was stained with heavy metals (rOTO protocol) and prepared using either critical point drying (CPD) or resin embedding (Epon). Imaging parameters include X-ray energy, effective pixel size, and exposure time.

| | sampleID | Paper Name | Staining | Embedding | Pillar diameter ($\mu\text{m}$) | Scan-Parameters | Figures |
| --- | --- | --- | --- | --- | --- | --- | --- |
| 1 | SK185_p1 | CPD1 | rOTO | CPD | 300 | E=33.6 keV; pixel size=100 nm; exposure time= 300 ms | Fig.4 |
| 2 | SK120 | CPD2 | rOTO | CPD | 300 | E=33.6 keV; pixel size=100 nm; exposure time= 300 ms | Fig.4 (averaged line-profile) |
| 3 | YY019_D1 | CPD3 | rOTO | CPD | 150 | E=33.6 keV; pixel size=60 nm; exposure time= 150 ms | Supp. Fig. S9 |
| 4 | SK186_p2 | Epon1 | rOTO | Epon | 300 | E=33.6 keV; pixel size=100 nm; exposure time= 300 ms | Fig.4 |
| 5 | C417F | Epon2 | rOTO | Epon | 300 | E=33.6 keV; pixel size=60 nm; exposure time= 330 ms | Supp. Fig. S10 |

**Figure 1.**
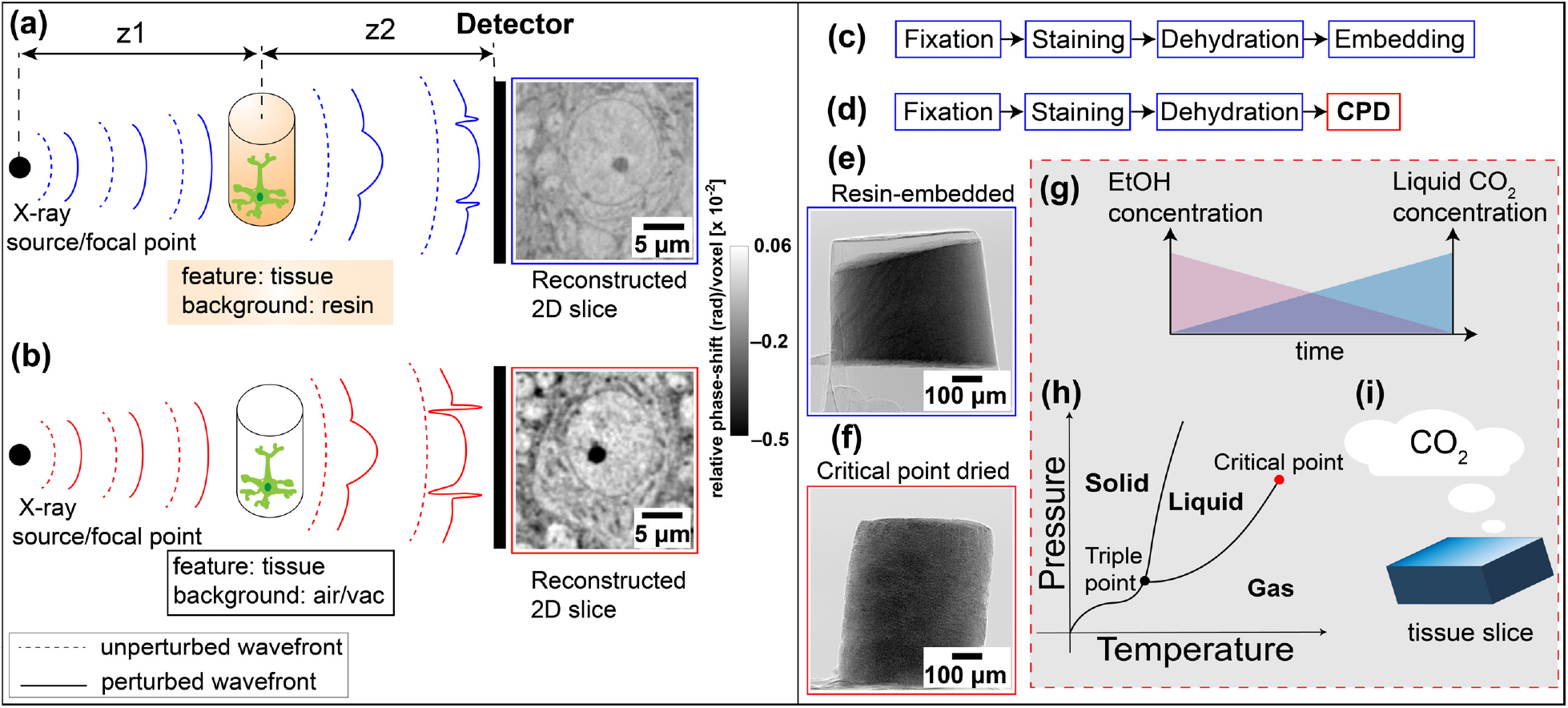
Sample preparation workflows and imaging principle for phase contrast imaging. **(a)** Schematic of X-ray propagation through resin-embedded, heavy metal-stained neuronal tissue. The difference in refractive index between tissue and surrounding resin causes modest distortion of the X-ray wavefront. z1 and z2 are the distances from the X-ray source or focal spot to the sample, and from the sample to the detector, respectively. **(b)** Equivalent schematic for critical point dried (CPD) tissue imaged in air or vacuum. The absence of embedding resin introduces a stronger refractive index discontinuity at the tissue–air interface, leading to enhanced wavefront distortion and increased phase contrast. **(c & d)** Standard preparation pipeline for Epon embedding **(c)** and CPD preparation **(d)**, both following fixation, staining, and dehydration. **(e & f)** Representative single-projection images of femtosecond-laser-milled cylindrical pillars imaged at 17 keV in air: Epon-embedded **(e)** and CPD-prepared **(f). (g-i)** CPD workflow: **(g)** ethanol-to-CO_2_ exchange gradient during CPD processing, **(h)** phase diagram of CO_2_ indicating the critical point transition, and **(i)** illustration of the final CPD step where liquid CO_2_ is brought to the critical point and then slowly vented as gas while maintaining temperature, avoiding surface tension forces associated with a liquid–gas interface and preserving ultrastructure [19].

We then assessed whether CPD would compromise the preservation of ultrastructure. We stained samples of mouse brain tissue, dehydrated them and dried them with CPD as described above. At that point, we infiltrated the dried sample with 100% resin. This provided a resin-embedded specimen fully compatible with transmission electron microscopy (TEM) (Fig. 2a). This approach allowed us to isolate the effects of CPD on the tissue while keeping fixation, staining, and dehydration protocols unaltered. Post CPD embedded samples displayed a preserved subcellular architecture (Fig. 2c), largely comparable to conventionally prepared samples (Fig. 2b). Qualitatively, in both the standard and post CPD embedded tissue, myelinated axons, mitochondria, nuclear envelopes, synaptic vesicle clusters, and fine neurites remained morphologically intact. A technical replicate of this experiment is provided in Supplementary Fig. S3, highlighting the reproducibility of this workflow across samples. These results confirm that CPD of a dehydrated biological soft tissue largely preserves the underlying ultrastructure.

**Figure 2.**
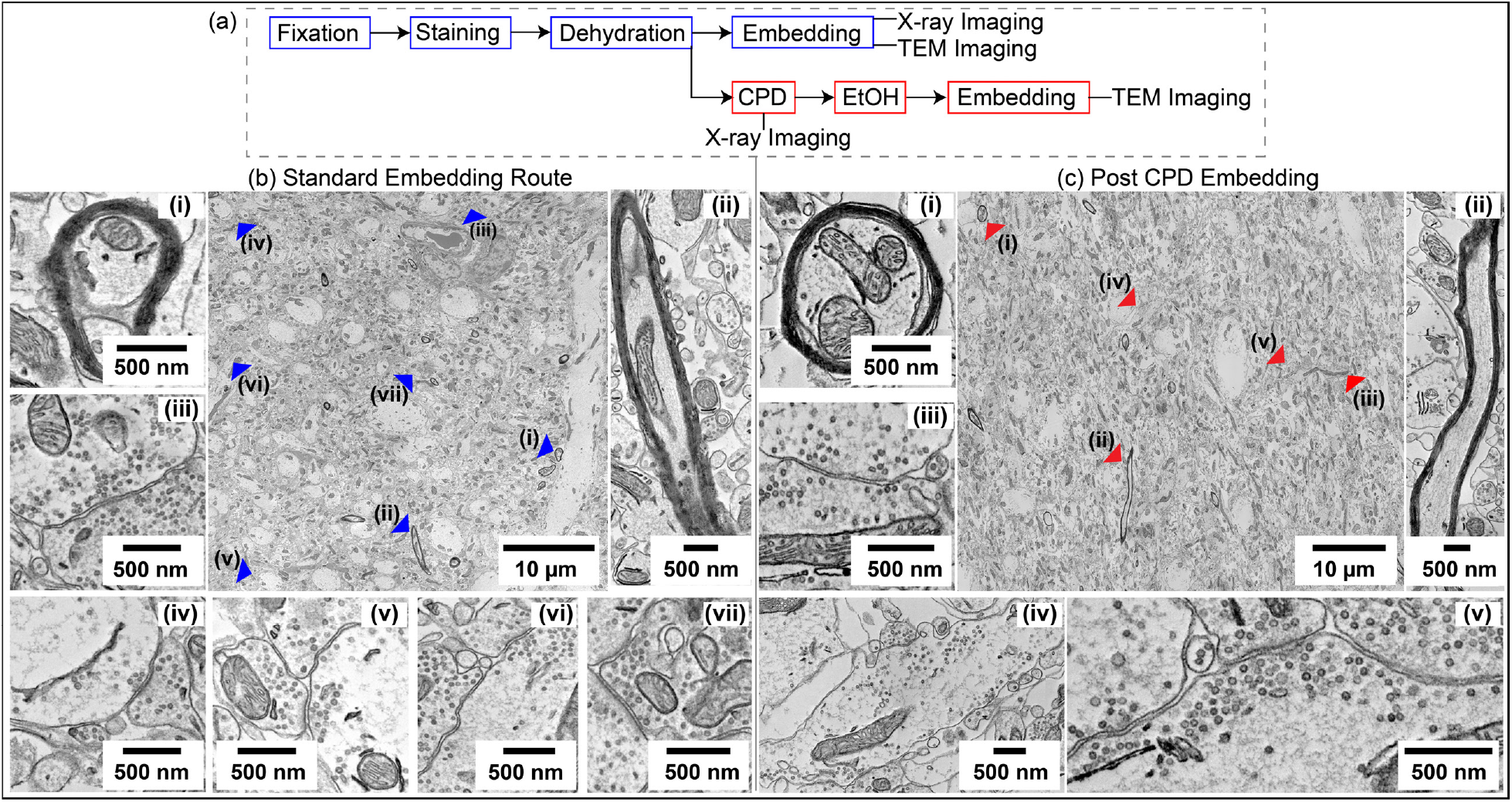
Ultrastructural preservation following critical point drying and post-CPD embedding in resin. **(a)** Divergent preparation workflows for TEM: tissue is either embedded directly in resin (top) or dried using CPD and subsequently embedded (bottom) prior to ultramicrotomy and TEM imaging. **(b)** TEM images of standard resin-embedded tissue show well-preserved ultrastructure, including clear nuclear membranes, myelinated axons, and synaptic vesicle populations. **(c)** CPD-treated tissue post embedded in resin retains qualitatively comparable morphological fidelity, with intact subcellular features throughout.

We next evaluated how CPD preparation influences image contrast in X-ray propagation-based phase contrast tomography (XPCT). We imaged (in air at an energy of 17 keV) CPD and resin-embedded samples at the EMBL High-Throughput Tomography (HiTT) setup of the P14 beamline (PETRA III), which employs a four-distance propagation-based acquisition scheme [22]. In both sample types, we could easily identify the biological features such as cell bodies, nucleoli, apical dendrites of projection neurons and glomeruli, which are commonly resolved in 325 nm effective pixel size XPCT datasets [2]. CPD samples showed however improved phase contrast in all those features (Fig. 3d). To quantify this effect we monitored signal in a subcellular feature that is widely distributed in all samples and that provides a consistently sharp contrast: the nucleolus. This subcellular compartment is located inside the nucleus, in the shape of a spheroid of approximately 2 μm in diameter in principal neurons [23] and is mainly composed of proteins and nucleic acids. Accordingly, like other protein clusters (such as postsynaptic densities) it accumulates heavy metals during the staining protocol. Importantly, its immediate surrounding, the nucleoplasm, has a much lower density of lipids and proteins, resulting in a sharply defined, compact and frequent feature that contains the limits of the dynamic range of signal in the dataset. Therefore, nucleoli, visible as small dark cores within the pale nucleus of mitral or tufted cells, the principle neurons of the mouse olfactory bulb, provided an excellent substrate for a comparative quantitative analysis of X-ray phase contrast signal.

**Figure 3.**
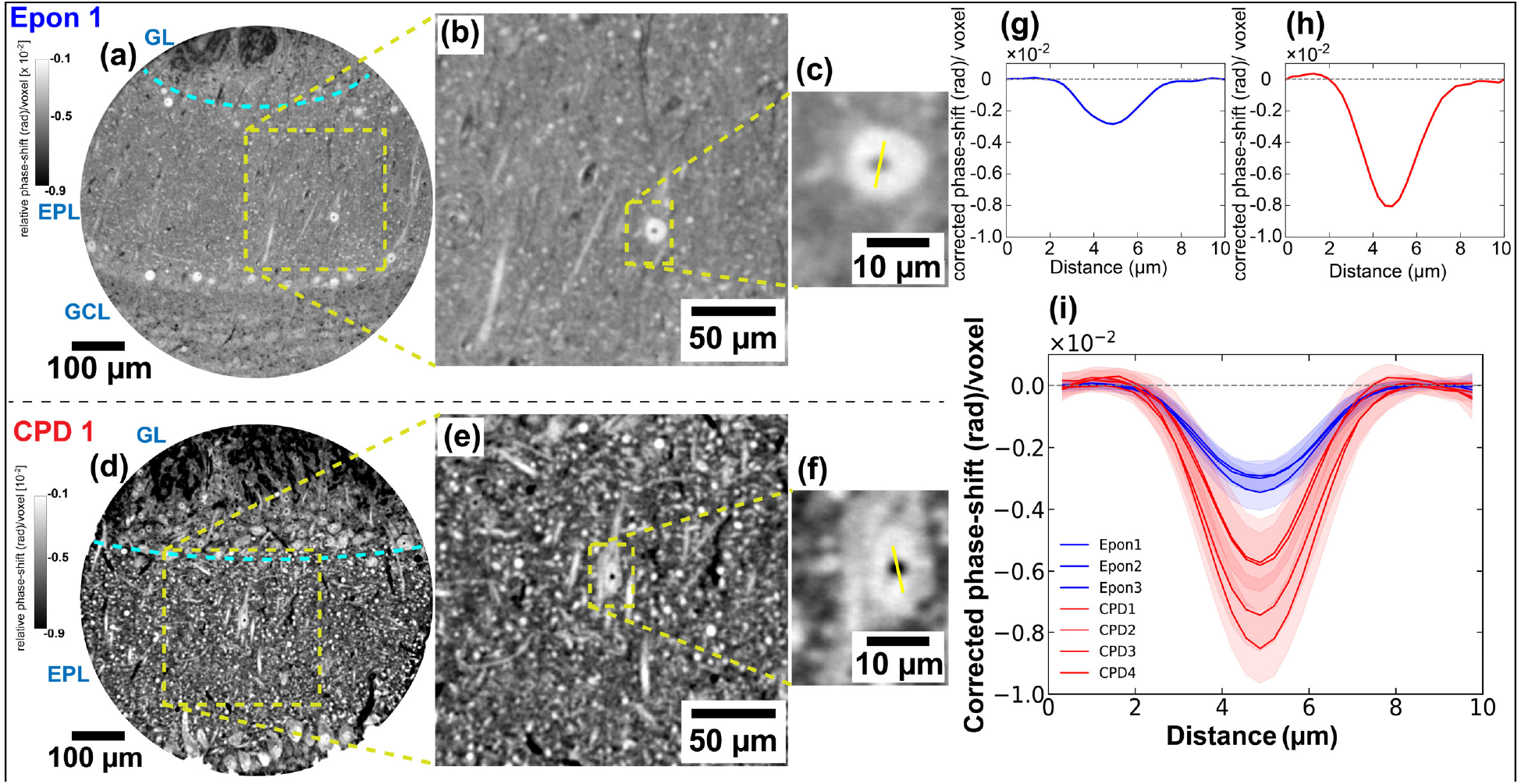
Enhanced phase contrast in CPD-treated tissue imaged via propagation-based XPCT at P14. **(a, d)** Reconstructed tomographic slices from embedded (Epon) and CPD-treated tissue pillars, respectively, centred on the external plexiform layer (EPL) of the olfactory bulb, and parts of the glomerular layer (GL) and granule cell layer (GLC) are also visible. **(b, e)** Magnified views within the EPL. **(c, f)** Showing nucleolus features (yellow line: profile axis), with corresponding corrected phase-shift profiles in **(g, h). (i)** Overlay of line profiles from multiple samples containing the nucleolus feature (Epon: blue, CPD: red), demonstrating consistent enhancement of phase-shift [corrected phase-shift = raw-signal − baseline mean (background)] in CPD samples. Shaded areas indicate standard deviation across measured regions. Reconstructed tomographic slices of embedded and CPD datasets labelled here are shown in Supplementary Fig. S4 and S5, respectively. Other individual line profiles from each dataset are shown in Supplementary Fig. S6. Increased phase contrast in CPD tissue results from the higher refractive index mismatch between tissue and surrounding air/vacuum, compared to resin-embedded preparations.

We extracted the image intensities of the voxels following a line that crossed the nucleolus across multiple nucleoli in each analysed tomogram (Fig. 3c, f) and corrected the signal obtained by subtracting the baseline measured in the adjacent nucleoplasm to robustly monitor the phase-shift across the nucleolus in CPD and Epon-embedded samples (Fig. 3g,h). CPD samples consistently exhibited deeper intensity troughs (Fig. 3g, h, i). The enhanced signal we observed in CPD samples reflects a stronger phase contrast triggered by ultrastructural features. Furthermore, a similar enhancement of X-ray scattering was also observed in unstained samples processed via CPD (i.e. aldehyde fixation is followed by dehydration without any staining step) when compared to unstained samples embedded in paraffin (Supplementary Fig. S7). Together, these results suggest that CPD preparation causes the enhancement in X-ray phase shift that leads to increased contrast in the images, independently of the staining method being used.

To assess whether the contrast improvements seen in CPD-prepared tissue extend into the nanoscale imaging regime, we next imaged CPD prepared samples with X-ray holographic nanotomography (XNH) at the ID16A beamline (ESRF). This imaging modality offers higher spatial resolution and increased coherence compared to propagation-based phase contrast. We configured scans using an isotropic voxel size of 100 nm, which in turn allowed obtaining single tiles of fields of view extending 300 μm in diameter. The ability to propagate wavefronts over extended distances in this configuration increases sensitivity to subtle phase gradients which enables the resolution of finer structural features. Imaging was performed under vacuum at 33.6 keV.

Both resin-embedded and CPD samples were obtained from a targeted and well-characterised anatomical region of the olfactory bulb, containing the external plexiform layer (EPL), as well as the adjacent glomerular layer (GL) (Fig. 4a, d). As in the P14 datasets, CPD samples display stronger phase contrast across both broad tissue interfaces and fine subcellular structures.

**Figure 4.**
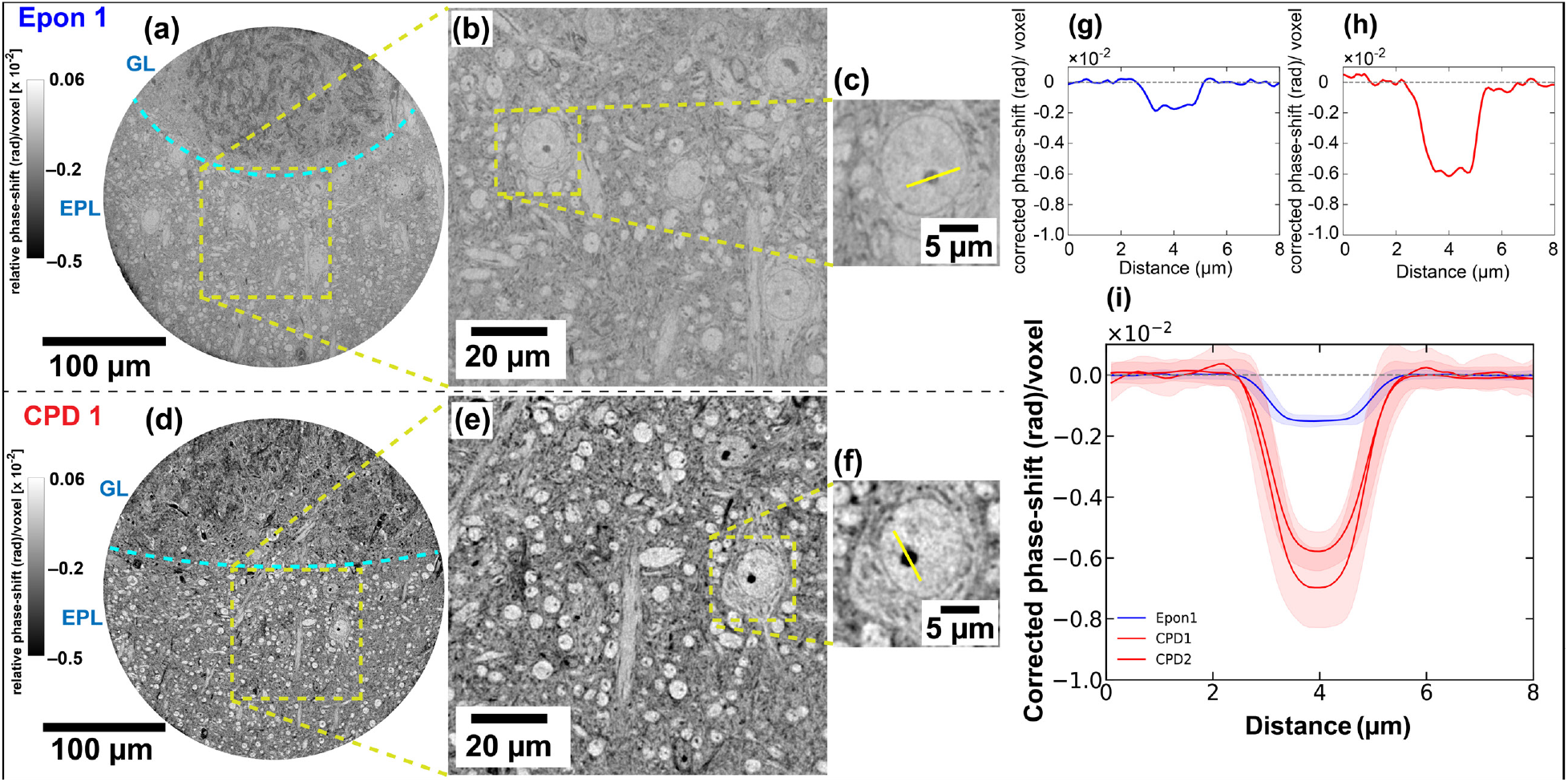
Enhanced phase contrast in CPD neuronal tissue imaged via XNH at ID16A. **(a, d)** Reconstructed slices from embedded (Epon) and CPD tissue, respectively, imaged in the external plexiform layer (EPL) and glomerular layer (GL) of the mouse olfactory bulb. **(b, e)** Magnified views within the EPL. **(c, f)** Showing nucleolus features (yellow line: profile axis), with corresponding corrected phase-shift profiles in **(g, h). (i)** Averaged line profiles from multiple nucleoli in Epon (blue) and CPD (red) samples, illustrating enhanced phase modulation [corrected phase-shift = raw-signal − baseline mean (background)] and gradient sharpness in dried tissue. Shaded bands represent standard deviation. Individual line profiles from each dataset is shown in Supplementary Fig. S8.

We quantified phase contrast in these datasets, as before, by characterising the signal amplitude across nucleoli (Fig. 4b-c, e-f). Line profiles drawn across these features (Fig. 4g, h) revealed an enhanced phase shift in CPD samples (Fig. 4i).

Overall, our findings show that CPD preparation of brain tissue samples introduces stronger X-ray scattering compared to standard resin-embedding protocols. CPD samples, devoid of any embedding material, exhibit a higher refractive index difference between foreground - lipids and proteins - and background - cytosol - increasing the contrast of tissue ultrastructure. We further compared the background-corrected phase shift amplitudes across all samples and beamlines explored (Fig. 5). CPD samples exhibit a 2-4*×* increase in phase contrast compared to their resin-embedded counterparts, an effect that can also be detected in unstained samples (Supplementary Fig. S7).

**Figure 5.**
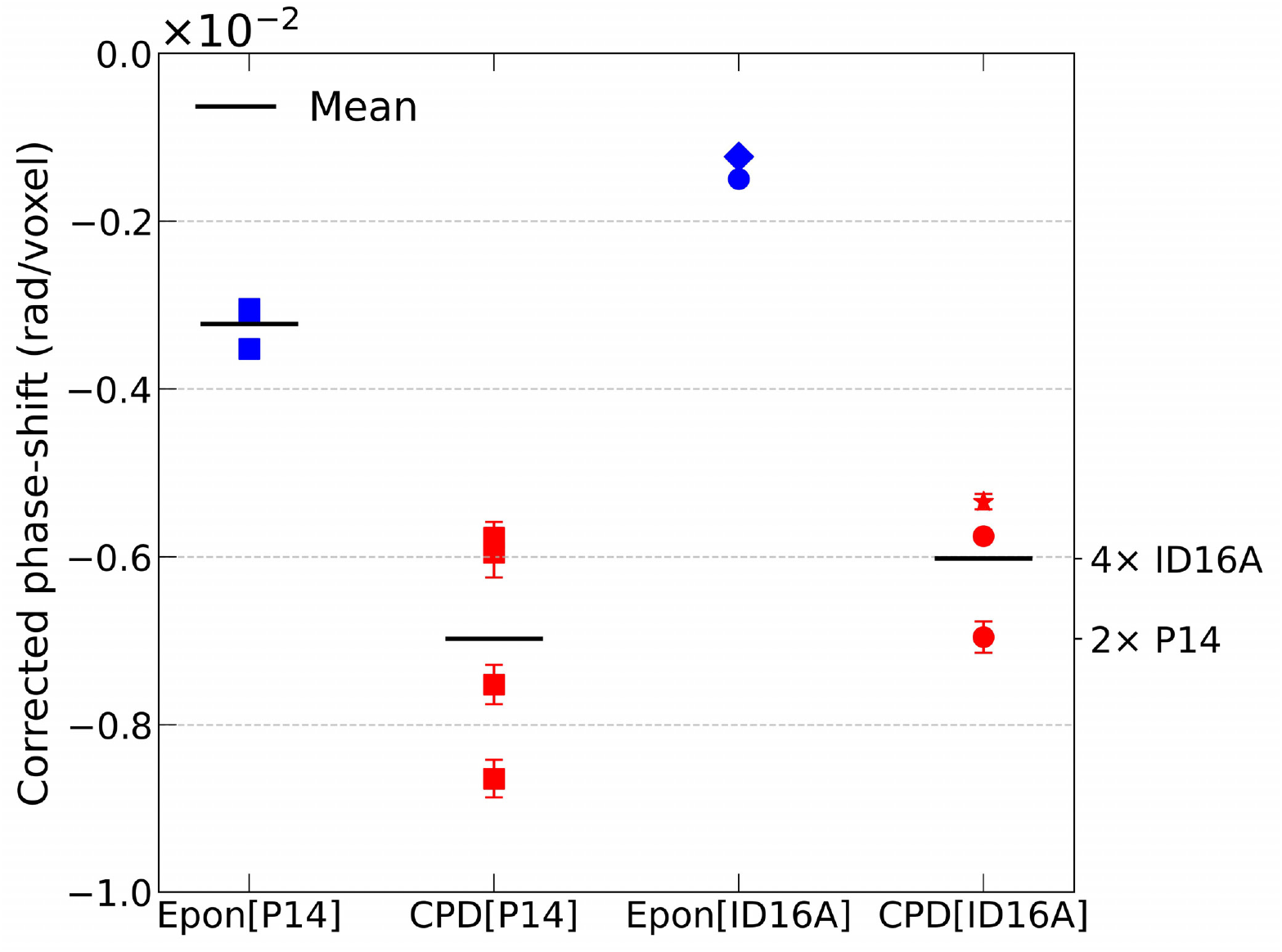
Summary of phase-shift enhancement across all datasets and imaging modalities. Amplitude of the corrected phase-shift (rad/voxel), extracted from averaged line profiles across nucleolar features in CPD-prepared (red) and Epon-embedded (blue) neuronal tissue. Data points include measurements acquired at P14 (squares) and ID16A (circles). Special cases (measured at ID16A) are marked: star indicates a different brain region (thalamus) and smaller pillar diameter (150 *µ*m), as shown in Supplementary Fig. S9; diamond indicates a dataset collected under different imaging conditions, as detailed in Supplementary Fig. S10. Despite differences in voxel size and imaging geometry, CPD consistently yields stronger phase-shift signals, confirming the generality of phase contrast enhancement introduced by CPD. Horizontal black bars indicate the mean value for each group.

## Conclusion

We present a method to prepare brain tissue samples optimised for X-ray phase contrast imaging. By converting tissue samples into a nanofoam-like structure, without any embedding material, the difference in scattering in subcellular and ultrastructural imaging is constrained between air/vacuum (little/no scattering), and the one provided by the membranes (which might be stained with heavy metals of choice).

We observe a 2-4*×* increase in background-corrected phase shift in high-signal biological features (nucleoli). This effect is consistent not only across samples but also across X-ray phase contrast imaging regimes and setups as well as in unstained samples, suggesting that it might apply as well to other X-ray phase contrast regimes such as X-ray ptychography.

This enhancement in phase contrast has meaningful implications for imaging efficiency. X-ray imaging dwell time depends quadratically on the inverse of the background-corrected signal amplitude [24]. Therefore, a 2-4*×* enhancement in signal should translate into a 4-16*×* speed-up in imaging. Hard X-ray imaging has demonstrated its capacity to resolve features at sub-10 nm detail [25] and ultrastructure in brain tissues below 40 nm detail [3, 9] non-destructively, making it an emerging technology suitable for scaling up connectomics to the whole mouse brain scale [26]. In this context, critical point drying (CPD) preparation of biological tissues can provide a boost in imaging speed, helping to bridge the current technological gap towards mm^3^-cm^3^ scale X-ray connectomics.

Sample ultrastructure remained largely preserved throughout CPD and follow-up resin embedding, as revealed by TEM, with neurites, synapses and vesicles displaying comparable patterns to the control specimens. Furthermore, femtosecond laser milling [21] provided a unique tool to manipulate the resulting metallic nanofoam samples, and to generate targeted pillars of controlled diameter (150-1000 μm). Together, this highlights a new technological framework for preparing tissue samples for ultrastructural imaging that carries multiple advantages. First, since this protocol eliminates the need of any resin infiltration step, it could bring a robust alternative to prepare samples of mm^3^-cm^3^ in volume [27, 28]. Second, this method unlocks the choice of resin from the X-ray phase contrast imaging step. Therefore, this enables CPD-prepared mm-scale tissues to be optimally imaged with X-rays before trimming them into arrays of 200 μm-wide samples with a femtosecond laser, which can then be individually embedded in the optimal resin for their targeted imaging endstation. Neighbouring tissue regions could therefore meet the standards to be imaged with follow-up Focussed Ion Beam SEM (by e.g. embedding that one pillar with durcupan) and with serial block-face SEM (by e.g. embedding that other pillar with Epon). And third, CPD sample preparation combined with whole mouse brain staining methods, femtosecond laser milling and high-throughput imaging approaches (such as those offered by high-throughput subcellular-resolution synchrotron X-ray phase contrast beamlines [22, 29–31]), offers a versatile and high-throughput sample preparation workflow that can help scale up the bandwidth of tissue life science nano-imaging experiments.

While this study focussed on improving background-corrected signal by reducing the background, further improvements may come from optimising staining protocols, particularly through the use of lighter elements with favourable X-ray scattering properties. These optimisations will likely provide further improvements in contrast and therefore in imaging speed, paving the way for tissue-scale X-ray connectomics. Moreover, optimised preparation of unstained biological tissues for X-ray imaging can broaden reach of tissue nanoimaging to scientific cases where heavy metal staining might pose a limitation. This advantage might therefore appeal to studies on unconventional animal species, exploring diverse developmental stages, or in locations where the logistics of heavy metal manipulation are difficult to implement - such as in fieldwork or in clinical setups aimed to explore volume ultrastructure of solid biopsies.

Finally, we report a versatile approach to create nanofoams with customised properties, using aldehyde-fixed biological tissues as a source material. These nanofoams can be tuned to match requirements by staining the fixed tissues with the metals of choice prior to dehydration. This approach is therefore compatible with optimising tissue staining to maximise X-ray phase contrast, but also to match requirements for other applications in which nanofoams might be of use.

Altogether, our findings show that removing interstitial material from biological tissues through CPD enhances X-ray phase contrast signal by 2-4*×*. This boost can in principle translate into a larger improvement in imaging efficiency, providing a means to scale-up X-ray phase contrast imaging workflows of biological tissues without compromising ultrastructural integrity.

## Methods

### Sample Preparation

All animal protocols were approved by the Ethics Committee of the board of the Francis Crick Institute and the United Kingdom Home Office under the Animals (Scientific Procedures) Act 1986.

A summary of all samples can be found in Tables 1 and 2.

#### Dissection & fixation

Mice were sacrificed and 300 - 500 *µ*m thick coronal brain sections containing the olfactory bulb (OB) region were cut using a Leica VT1200S vibratome. Slicing was performed in an ice-cold dissection solution containing 65 mM NaH_2_PO_4_·H_2_O (phosphate buffer), 4.6% (146 mM) sucrose, 0.6 mM CaCl_2_,, and 0.02% sodium azide (adjusted to an osmolarity of 300 *±* 20 mOsm/L), bubbled with 95% O_2_ /5% CO_2_ to adjust pH, to keep the tissue moistened and cold throughout dissection and slicing. Immediately following sectioning, tissue was transferred into ice-cold fixative, consisting of a mixture of 1.25% glutaraldehyde and 2.5% paraformaldehyde in 150 mM sodium cacodylate buffer (pH 7.40, 300 ± 20 mOsm/L). Samples were incubated in fixative overnight at 4°C. The next day, tissues were washed three times for 10 minutes each in wash buffer (150 mM sodium cacodylate, pH 7.40, 300 *±* 20 mOsm/L) at 4°C. All steps were carried out in ice-cold, osmolarity-verified buffers to ensure tissue preservation.

#### Staining

Samples were processed using a standard heavy metal protocol (rOTO) [4]. Tissue was incubated in 2% osmium tetroxide (OsO_4_) in 0.15 M sodium cacodylate buffer (NCB) for 1.5 hours at 20°C, followed immediately by 2.5% potassium ferrocyanide in 0.15 M NCB pH 7.40 for an additional 1.5 hours at 20°C, without an intermediate water wash. Samples were then treated with 1% thiocarbohydrazide (aqueous) for 45 minutes at 30°C, followed by a second osmication step using 2% OsO_4_ (aqueous) for 3 hours at 20°C. Afterwards, samples were incubated in 1% uranyl acetate (aqueous) for an overnight at 4°C and later warmed to 50°C for 2 hours. Lead aspartate staining (aqueous, pH 5.0) was performed for 2 hours at 50°C. Water washes were carried out between each staining step unless stated otherwise.

#### Resin embedding

Following staining, samples were dehydrated through a graded ethanol series (75%, 90%, 100%, 100%), transitioned into propylene oxide. After this step, one set of samples underwent infiltration with increasing concentrations of hard Epon resin [Epon812 (TAAB T023, DDSA (TAAB D026), MNA (TAAB M011) and BDMA (TAAB B008), mixed as described in ([3, 32])] diluted in propylene oxide (25%, 50%, 75%, 100%, 100%). Resin-infiltrated samples were polymerised for 72 hours at 60–70°C. One of these samples (C417Fb) was embedded in a similar resin that used a different epoxy, EMbed812 (E.M.S. 14900).

#### Critical point drying

Critical point drying (CPD) followed directly after ethanol dehydration. CPD was employed to preserve tissue morphology by avoiding the damaging effects of surface tension that arise during conventional air drying. In hydrated biological specimens, evaporation at the liquid–air interface can introduce substantial tangential forces, leading to deformation, collapse, or delamination of micro- and nanoscale structures [15–18]. CPD circumvents this by transitioning from the liquid to the gaseous phase without crossing a distinct phase boundary. Because the critical point of water (374°C, 229 bar) is incompatible with biological preservation, water was first replaced with a series of intermediate exchange solvents, ethanol in this case, which is miscible with both water and liquid CO_2_. Ethanol was then gradually exchanged with liquid CO_2_. Unlike ethanol or acetone, which require extreme conditions for CPD (critical points above 230°C), CO_2_ reaches its critical point at 35°C and 70 bar, making it suitable for biological applications. Once in the CPD chamber (Leica EM CPD300), the liquid CO_2_ was brought to its critical point and slowly vented by reducing the pressure while maintaining the critical temperature, thus drying the sample minimising phase transition forces. The parameters used for CPD on the Leica EM CPD300 were as follows: CO_2_ In (speed = slow, fillers = full, delay = 120 seconds), Exchange (speed = 5, cycles = 14) and Gas Out (heat = medium, speed = slow /100%), resulting in a total process duration of approximately 1 hour and 40 minutes.

#### Resin embedding after CPD

Dried samples were incubated with 100% ethanol for 4 hours at 4°C (changing ethanol at every hour mark), and polymerised following the standard resin embedding procedure.

#### Femtosecond laser milling

Both sets of resin embedded and dried tissue samples were then milled to cylindrical shapes with a femtosecond laser (Optec Femtosecond laser WS Starter equipped with a Coherent Monaco laser with dual wavelength, 515 nm and 1030 nm; all pillars in this study are milled with wavelength = 515 nm). The diameters of the final cylinders were decided so samples would fit in the field of view of the tomography experiments at synchrotron beamlines. Pillars with diameter ≈ 500 *µ*m and ≈ 300 *µ*m were milled for measurements at P14 and ID16A, respectively (Figure 1 & Supplementary Fig. S2).

### Transmission Electron Microscopy

To confirm ultrastructure preservation, transmission electron microscopy (TEM) experiments were performed on thin-sections of ≈ 80 nm thickness (obtained via Ultramicrotome sectioning) in both resin embedded and post CPD embedded samples. JEOL 1400FLASH electron microscope was used for this purpose. Beam current was set to 120 kV and imaging was performed at 12k magnification (giving a pixel size of 1.4 nm), and JEOL LLQ software suite was used for image acquisition and stitching to cover a wide region of interest (Figure 2, Supplementary Fig. S3).

### Plasma-Focused Ion Beam Milling

Plasma FIB milling was carried out on a Thermofisher Scientific Helios Hydra using 30 kV Xe^+^ ions which has a current range between 1 pA and 2.5 *µ*A. SEM imaging was carried out using using the Ion Conversion Electron Detector (ICE) in secondary electron detection mode (Supplementary Fig. S1 (a) 10 kV, 100 pA, (c) 2 kV, 100 pA, (e) 2 kV, 100 pA), the Through Lens Detector (TLD) in Field Free secondary electron mode (Supplementary Fig. S1 (b) 2kV, 100 pA), and the Through Lens Detector (TLD) in Immersion mode with backscatter electron mode (Supplementary Fig. S1 (d) 2 kV, 100 pA). Cross-sectional milling at 30 kV was performed using a range of beam currents (60, 200, 500 and 1000 nA) to identify the fastest approach for achieving a relatively undamaged surface finish. It was found that 500 nA was sufficient to quickly mill the cross sections shown in Supplementary Fig. S1 (b) and (c). 12 kV Xe^+^ ions were used to deposit a 10 um × 10 um × 1 um Pt/C protective layer over a region of interest to protect the surface from milling during the shape of a pillar structure.

### X-ray phase contrast tomography at P14

X-ray phase contrast tomography was performed at the High-Throughput Tomography (HiTT) endstation of the EMBL beamline P14 at PETRA III (DESY), optimized for in-line phase contrast imaging of mm-sized biological samples. Brain tissue pillars were vertically mounted on custom 3D printed holders and scanned at ambient pressure and room temperature. Each tomographic scan used four sample-to-detector distances to enable multi-distance phase retrieval [5]. All experiments at P14 were performed at an X-ray energy of 17 keV. A high-resolution detection system (Optique Peter) equipped with an LSO:Tb on YbSO scintillator (thickness 8-10 *µ*m) and a 20*×* magnifying microscope objective was coupled with a PCOedge 4.2 sCMOS camera (2048 *×* 2048 px, 6.5 *µ*m pixel size, max. 100 Hz frame rate). This resulted in an effective pixel size of 0.325 *µ*m, yielding a field of view of 666 *×* 666 *µ*m^2^, which was sufficient to cover the full pillar diameter (500 *µ*m) in a single scan. The acquisition covered 181° of rotation with 1800 projection angles. The raw intensity projections at each distance were flat-field corrected, aligned, and subsequently processed using a non-iterative, contrast transfer function (CTF)-based phase retrieval algorithm (where the refractive index decrement δ and absorption index *β* were set to that of osmium, and *δ/β* value of 10 was approximated for X-ray energy of 17 keV) [5]. After retrieval, projections were subjected to tomographic reconstruction using grid-rec algorithm to obtain volumetric datasets. The HiTT pipeline also includes on-the-fly data reduction and speed-optimized multi-node phase retrieval and tomographic reconstruction, which allows for real-time feedback of reconstructed tomogram images to image acquisition parameters, enabling an efficient handling of multiple samples [22].

### X-ray holographic nanotomography at ID16A

X-ray holographic nanotomography (XNH) was performed at the ID16A beamline of the European Synchrotron (ESRF, Grenoble, France). The beamline’s endstation is located at 185 m from the source to improve spatial coherence and full-field nanoscale imaging is achieved thanks to nanofocussing with multilayer-coated Kirkpatrick-Baez mirrors [33]. We imaged at 33.6 keV with a spot size of around 15 nm. Sample pillars were mounted on a high-precision rotation stage within a vacuum chamber maintained at 10^−7^ mbar. The detector was placed outside the vacuum chamber at about 1.289 m from the focal spot. For each complete acquisition, we acquired four tomographic scans over 180 deg (2000 projections each), at different propagation distances. For each rotation angle, the four corresponding holograms were flat-field corrected, brought to the same magnification and aligned in order to combine them for obtaining a phase map. Phase retrieval was then carried out using an iterative multi-distance approach, starting from an approximation regularized with a δ*/β* ratio of 27 (corresponding to Os at 33.6 keV) [34], and only the phase term was updated per iteration while keeping the amplitude constant. Retrieved phase maps were used for tomographic reconstruction using a filtered back-projection algorithm to generate isotropic 3D volumes at 100 nm voxel size (for all datasets). The resulting datasets allowed for cellular feature discrimination across entire pillar volumes [1, 4, 9, 35].

### Data analysis

We used Fiji [36] to obtain gray values from voxels at regions of interest. Retreived data was then structured, analysed and plotted using custom scripts in python. All reconstructed datasets were then stored in OME-ZARR format into a webknossos environment (scalable minds, [37]) for data sharing purposes (see Data and Code Availability sections).

## Acknowledgments

We thank Kevin Briggman and Thomas Kuner for inspiration, critical insights and discussion on the CPD protocol. The authors are grateful to the biological research, scientific computing and electron microscopy science technology platforms of the Francis Crick Institute (London, UK). We thank the Bionanofabrication clean room at the Imperial College (London, UK) for assistance in the operation of the femtosecond laser system, and to Chiara Licusati & George Konstantinou (Francis Crick Institute) for early support in the operation of the femtosecond laser system. We thank Gleb Bourenkov for assistance during synchrotron beamtimes at P14 and William Chevremont for assistance at ID16A. We acknowledge the DESY, Hamburg, Germany and ESRF, Grenoble, France for provision of synchrotron radiation beamtime at the P14 EMBL beamline of DESY (proposal XIMG-13) and at the ID16A beamline of the European Synchrotron (proposals IHMA106 and LS3231). Access to the Thermofisher Helios Hydra system (plasma-FIB) was supported by funding from the EPSRC (EP/V007661/1 to J.D.). For the purpose of Open Access, the author has applied a CC BY public copyright licence to any Author Accepted Manuscript version arising from this submission. This work was supported by the Francis Crick Institute, which receives its core funding from Cancer Research UK (CC2036 to A.T.S.), the UK Medical Research Council (CC2036 to A.T.S.), and the Wellcome Trust (CC2036 and 110174/Z/15/Z to A.T.S.). It was also supported by a Physics of Life grant (EP/W024292/1) to A.T.S. and A.P. funded by EPSRC and Wellcome and by the IMAGINE grant from the European Commission (101094250) to L.D., P.C. and A.P. A.P. acknowledges funding from the European Research Council under the European Union’s Horizon 2020 Research and Innovation Programme (852455).

## Author Contribution Statement

Conceptualization: SK, ATS, CB. Sample preparation: SK, YZ, YY, FS, JOD. XPCT data acquisition at P14: SK, JA, YZ, JR, AS, FM, KD, YY, ED, CB. XNH data acquisition at ID16A: SK, AV, YZ, JR, YY, PC, AP, ATS, CB. TEM data acquisition: SK. Data analysis: SK. Manuscript first draft: SK, CB. Figure preparation: SK, CB. All authors contributed to the editing of the manuscript.

## Data Availability

All 3D reconstructed tomograms reported in this study and tables containing source data for all quantitative analyses are accessible through the associated code repository (see Code Availability).

## Code Availability

Analysis code is available from https://github.com/safekhan/CPDtissuesXray.

## Supporting Information

**Figure S1.**
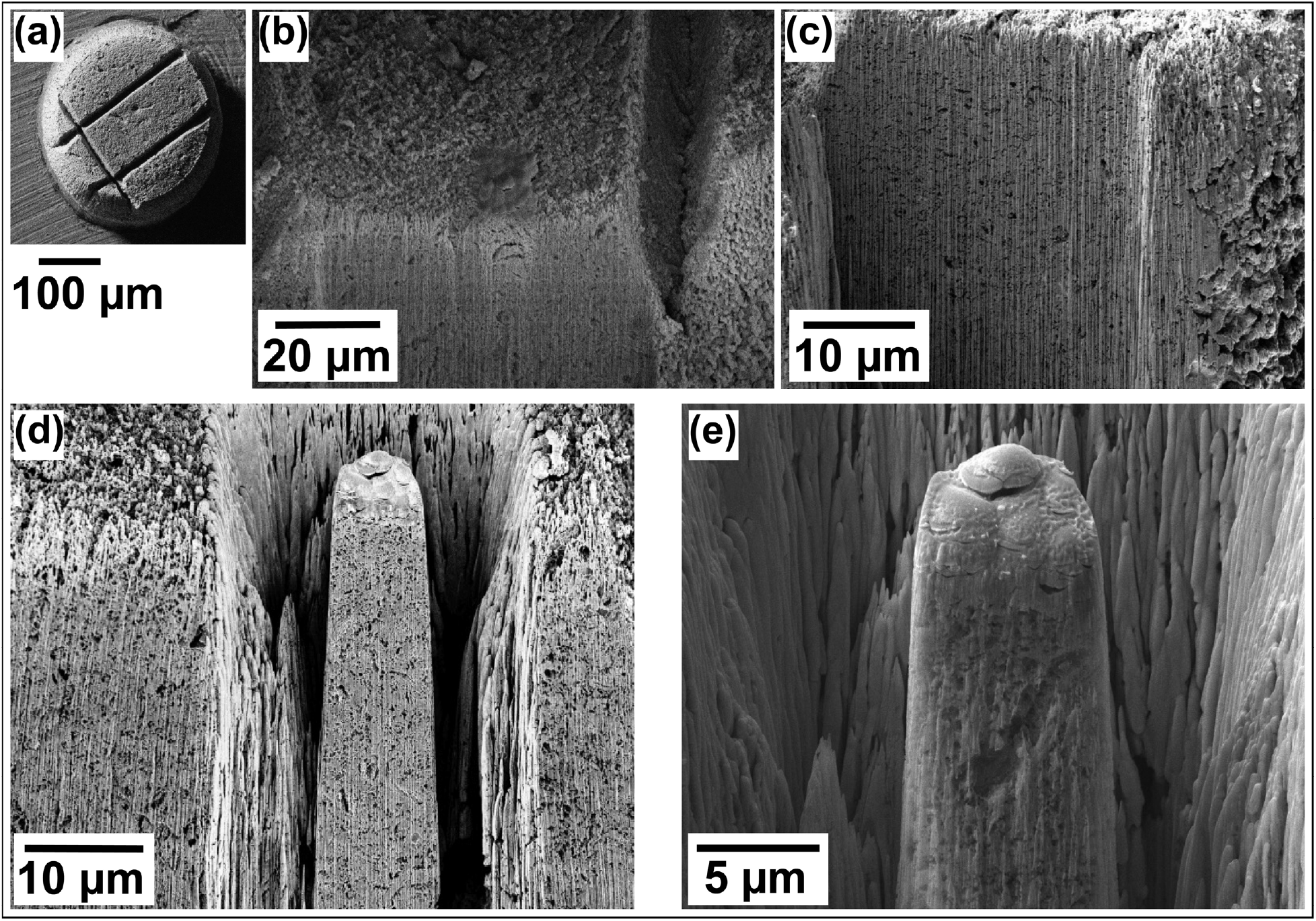
Plasma-Focused Ion Beam (p-FIB) milling of a CPD-prepared, metal-stained brain tissue sample reveals porous, nanofoam-like structure. **(a)** Top view SEM image of a ≈ 300 μm diameter CPD pillar prepared by femtosecond laser milling from a CPD-treated sample. **(b-c)** Higher magnification SEM images of the same pillar after targeted edge milling using xenon plasma-focused ion beam (p-FIB). The striated and porous appearance across the milled cross-sections reflects the air-filled nanostructure characteristic of a metallic nanofoam, consistent with the absence of embedding material. **(d-e)** On the same CPD pillar described in (a-c) we sculpted a smaller ≈ 10 μm diameter pillar using p-FIB milling.

**Figure S2.**
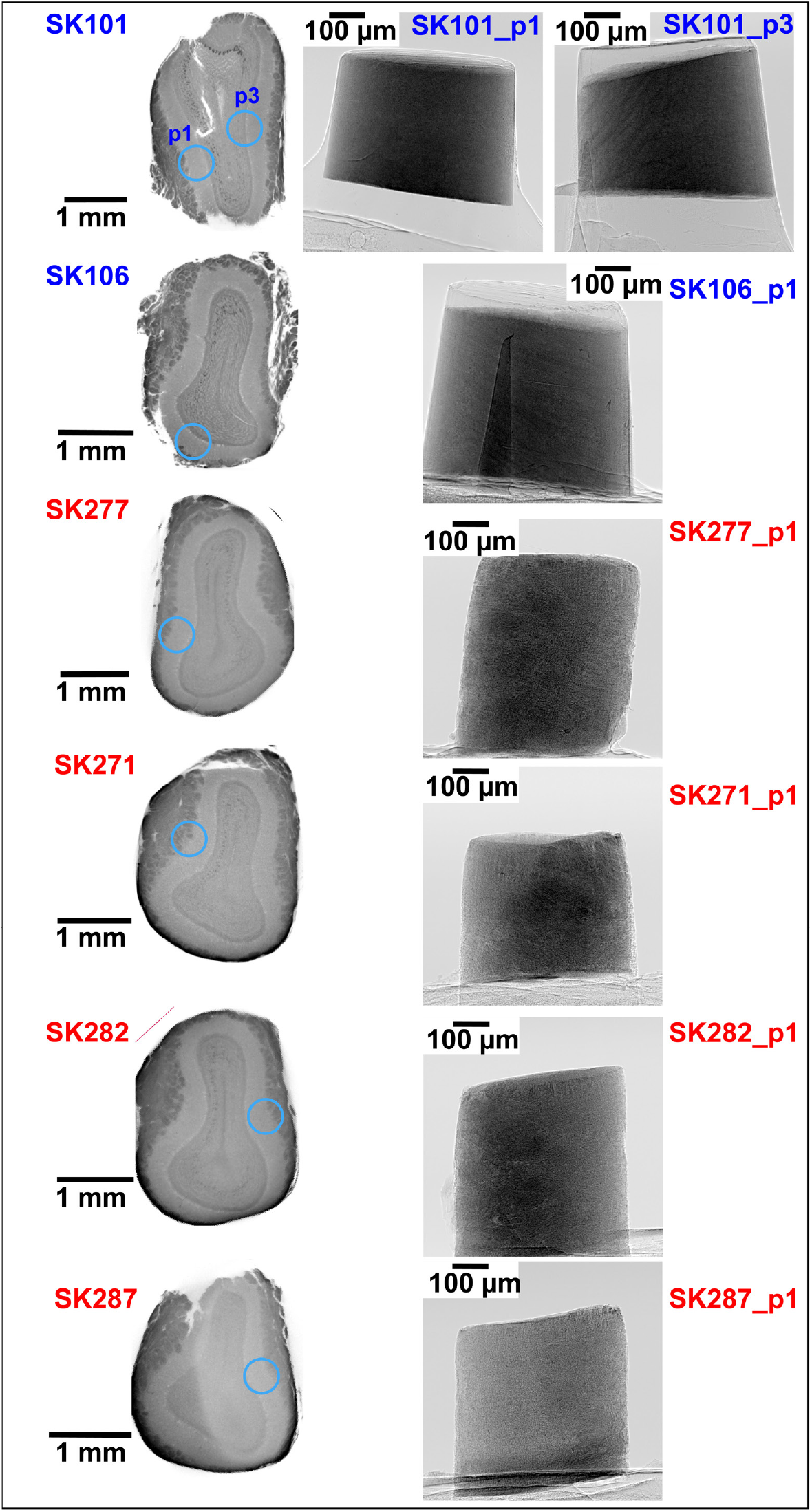
Overview of pillar locations for all samples imaged at P14. **(a)** Lab-based X-ray micro-CT scans (left) show coronal views of tissue slices from which pillars were extracted using femtosecond laser milling. Blue circles indicate the targeted regions where pillars were milled. Each slice corresponds to either resin-embedded (SK101, SK106; blue labels) or CPD-prepared (SK277, SK271, SK282, SK287; red labels) tissue. Corresponding single-projection images (right) acquired at P14 highlight pillar morphology and mounting geometry for each sample.

**Figure S3.**
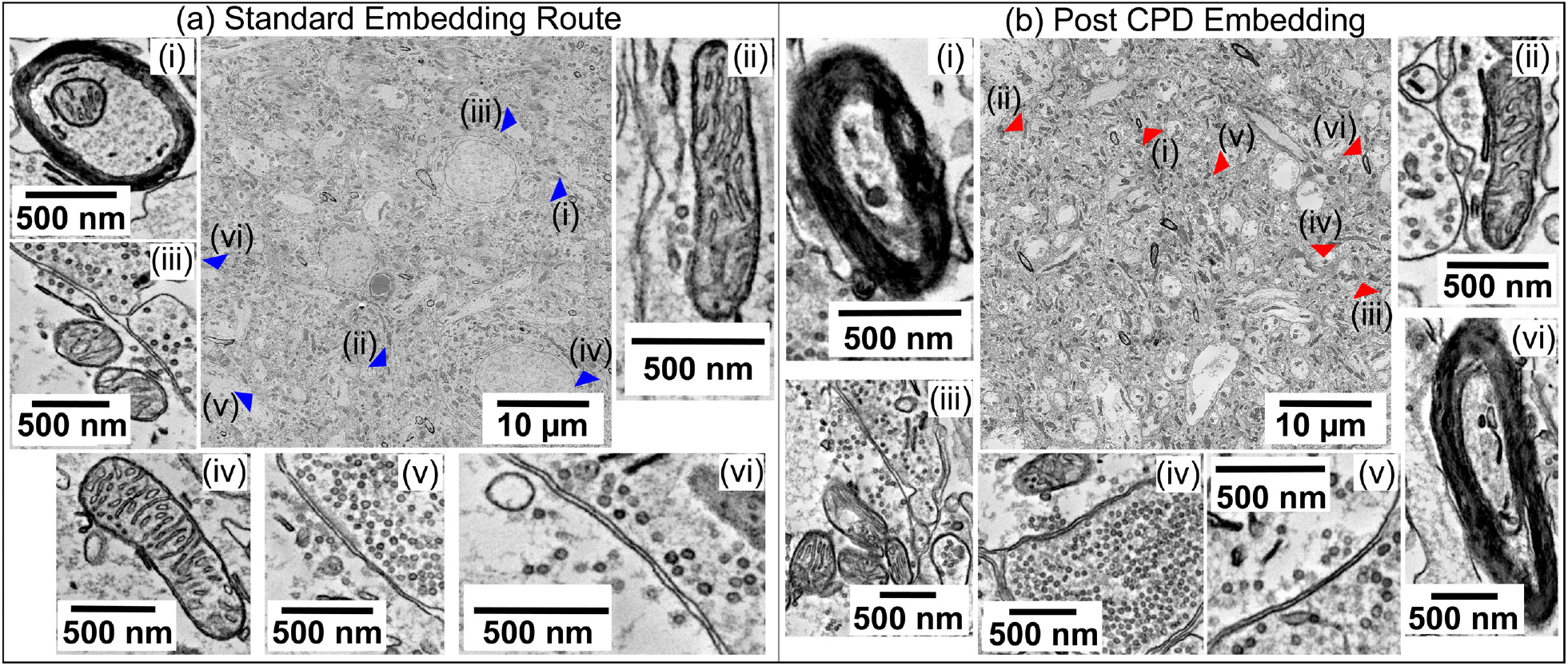
Consistent ultrastructural preservation in post-CPD embedded tissue across independent samples. Transmission electron microscopy (TEM) images from resin-embedded **(a)** and post CPD embedded tissue **(b)**, prepared from separately processed brain sections. Across both conditions, qualitatively, key cellular features such as mitochondria, vesicle pools, nuclear membranes, and synaptic boutons remain structurally intact.

**Figure S4.**
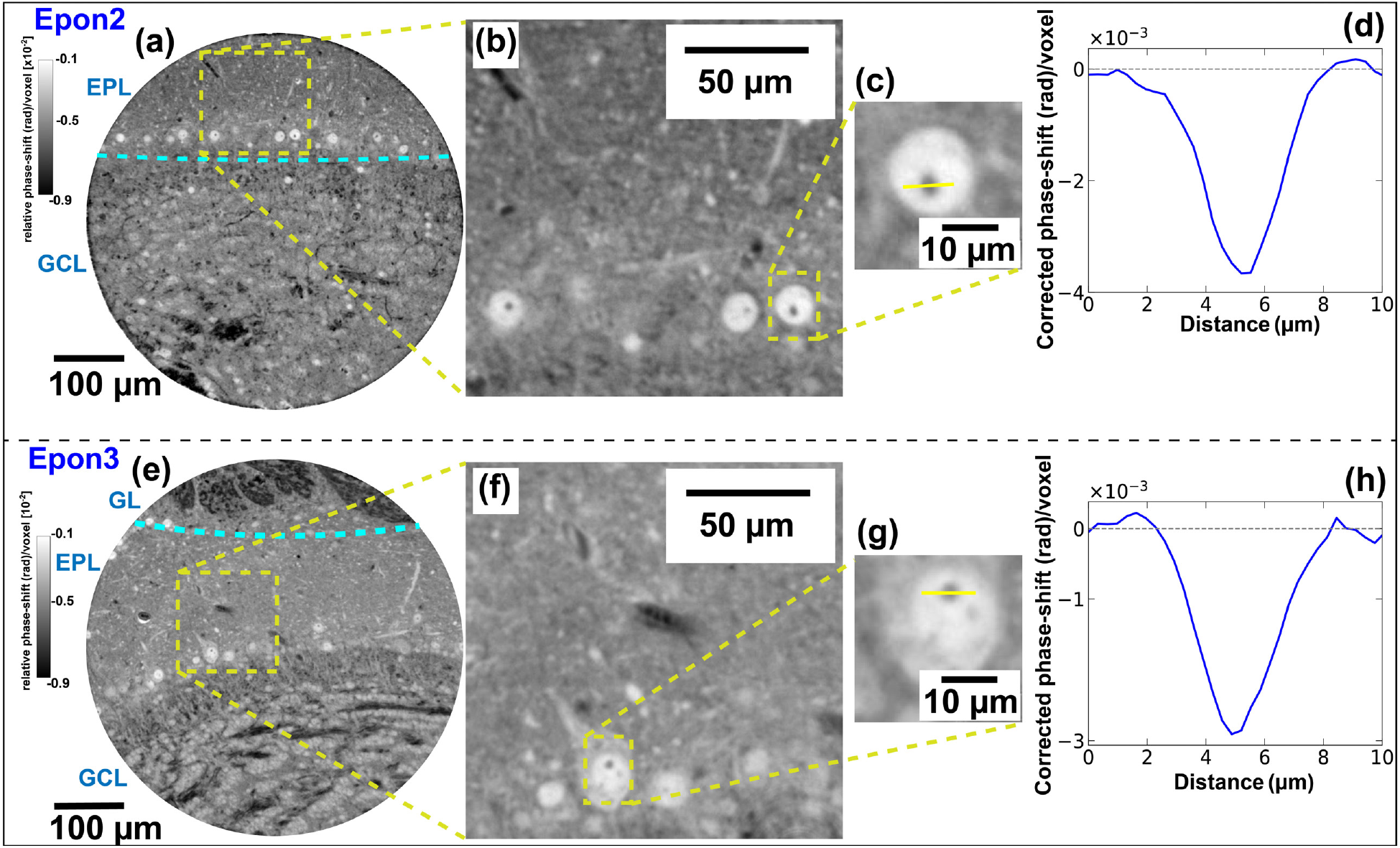
Examples of reconstructed slices of additional Epon-embedded samples imaged at P14 using X-ray phase contrast tomography. **(a-d)** Image and representative signal trace of sample labelled as Epon2 in the Fig 3 (i). **(e-h)** Image and representative signal trace of sample labelled as Epon3 in the Fig 3 (i).

**Figure S5.**
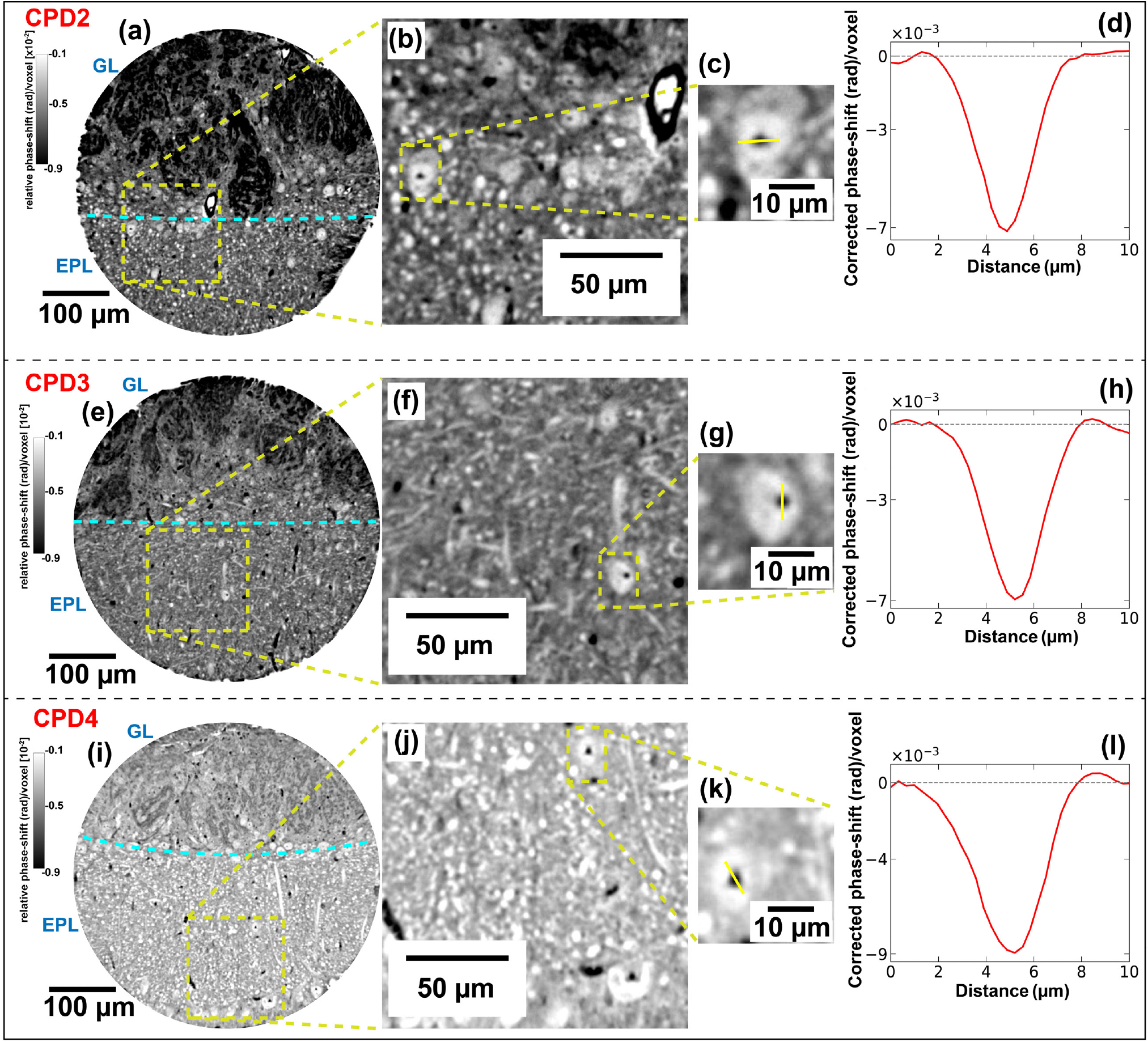
Examples of reconstructed slices of additional CPD samples imaged at P14 using X-ray phase contrast tomography. **(a-d), (e-h) & (i-l)** are images and representative signal traces of samples labelled as CPD2, CPD3, CPD4 in Fig 3 (i) in the main text.

**Figure S6.**
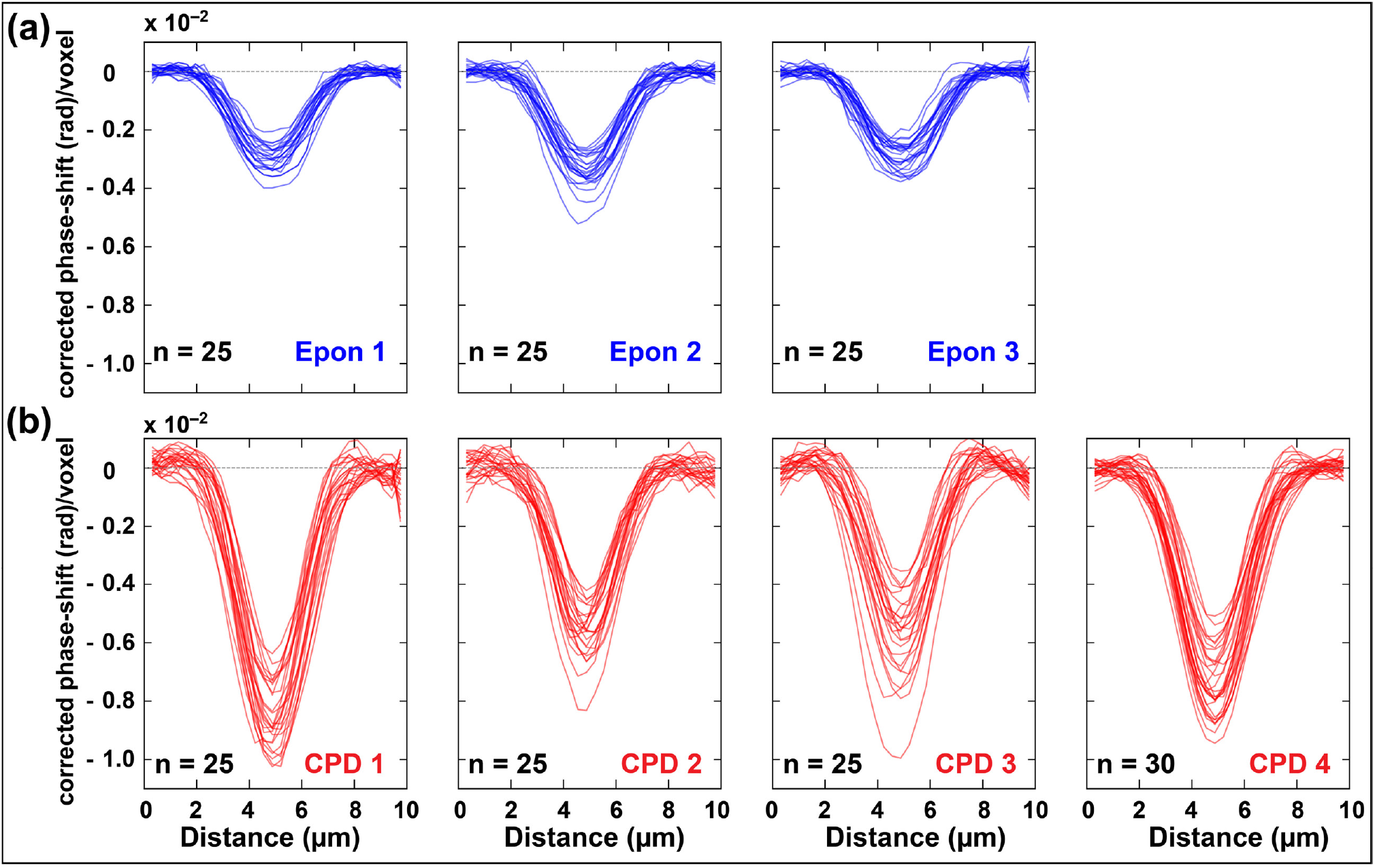
Background corrected line profiles for all the samples imaged at P14 using X-ray phase contrast tomography. Corrected phase-shift = raw signal − baseline mean (background). Line profiles were extracted across nucleoli (where n = number of line profiles) from multiple Epon **(a)** and CPD **(b)** datasets. The averaged line profiles from these datasets are shown in Figure 3i.

**Figure S7.**
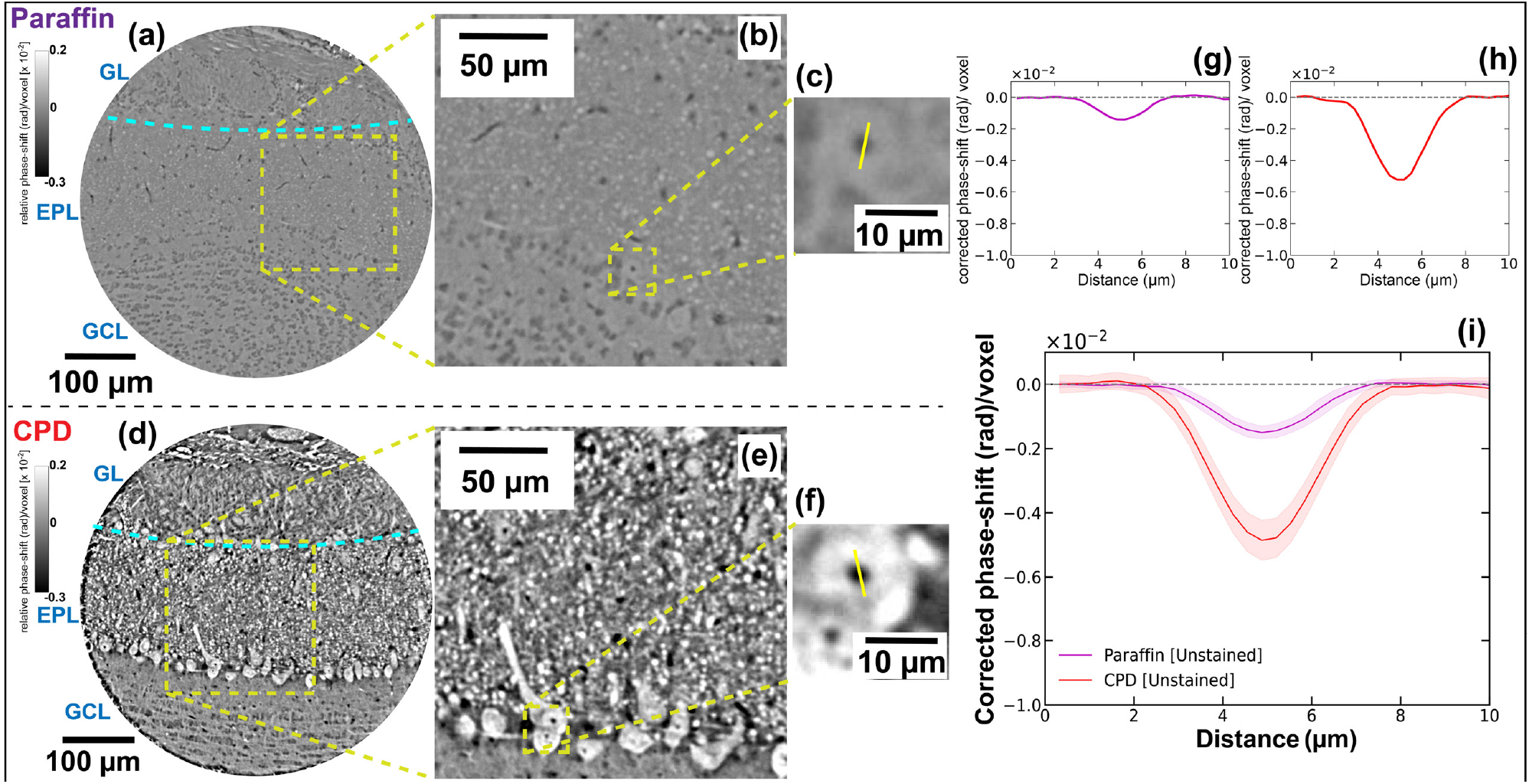
X-ray phase contrast signal in unstained tissue, CPD vs paraffin-embedded. Tomographic slices from mouse brain olfactory bulb tissue samples prepared without any staining and processed via either CPD or standard paraffin embedding, measured using XPCT at P14. **(a–c)** Paraffin-embedded sample. **(d–f)** CPD-prepared sample. **(g-h)**Representative line profiles in both samples. Mean *±* s.d. of multiple (n=25) measurements in this dataset.

**Figure S8.**
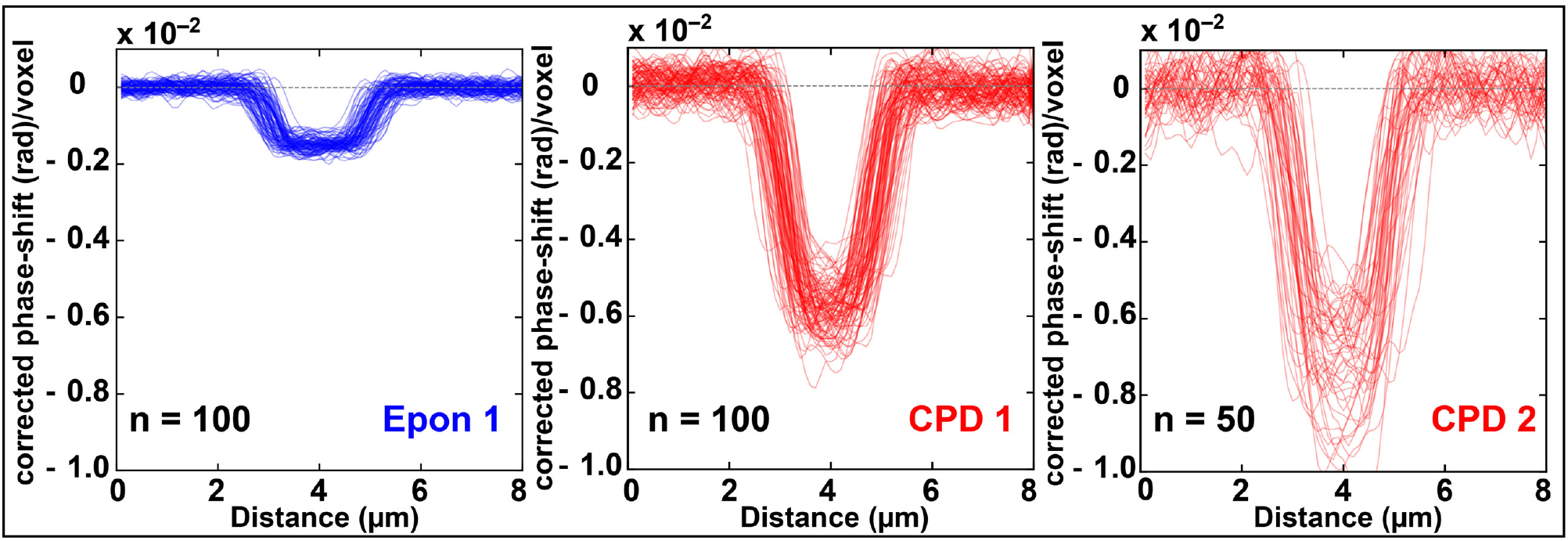
Background corrected line profiles for all the samples imaged at ID16A using X-ray nano-holotomography. Corrected phase-shift = raw signal − baseline mean (background). Line profiles were extracted across nucleoli (where n = number of line profiles) from Epon (blue) and CPD (red) datasets. The averaged line profiles from these datasets are shown in Figure 4i.

**Figure S9.**
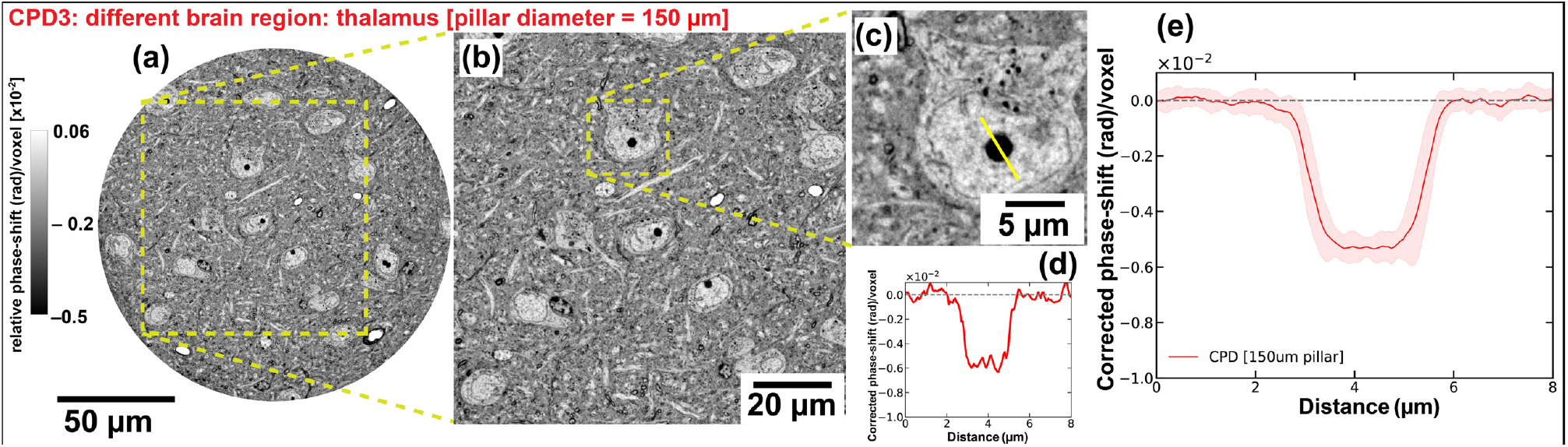
XNH of CPD-prepared sample from a different brain region (thalamus) and imaging configuration. **(a)** Reconstructed phase slice from a CPD-prepared sample imaged at ID16A (ESRF) from the thalamic region, using a 150 *µ*m diameter pillar. The voxel size was 60 nm, and exposure time was 150 ms. **(b-c)** Zoom-in views reveal preserved ultrastructure, including cellular and subcellular compartments. **(d)** Line profile extracted across a nucleolus (yellow line in c), showing strong phase contrast at the nuclear boundary. **(a)** Mean *±* s.d. of multiple (n=25) measurements in this dataset. This dataset contributes one of the CPD data points in Figure 5.

**Figure S10.**
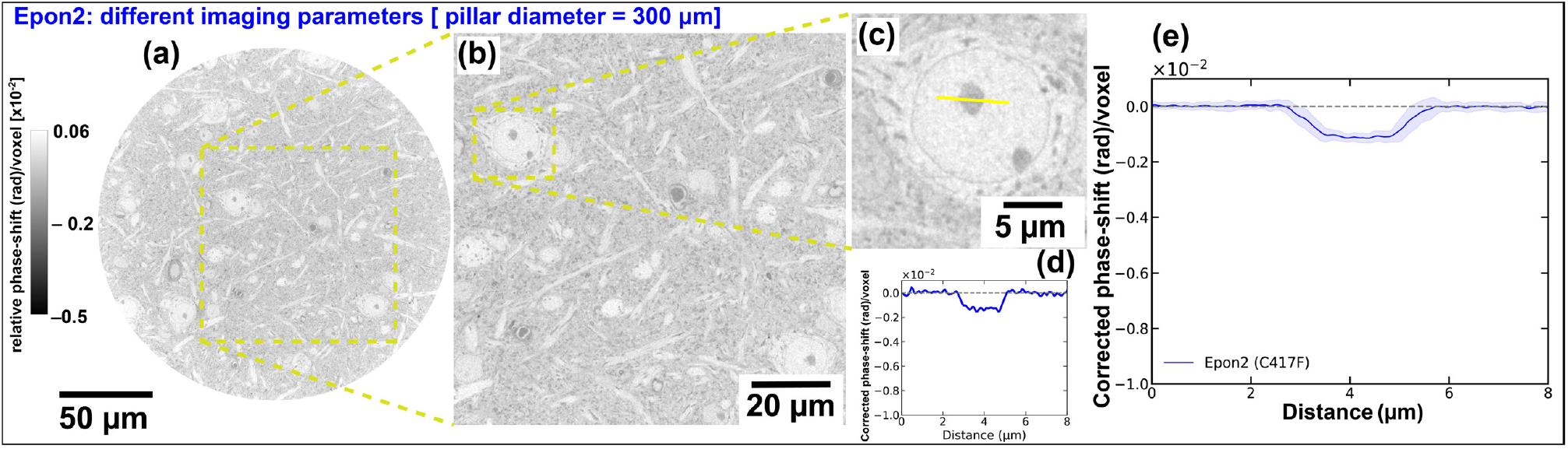
XNH of resin-embedded tissue (C417F) imaged with different parameters at ID16A. **(a)** Reconstructed phase slice of an Epon-embedded olfactory bulb sample imaged at ID16A. This scan was acquired with a 60 nm pixel size. **(b-c)** Zoom-in views of the same region highlight subcellular structures including cell somata and nucleoli. **(d)** A representative line profile across a nucleolus (yellow line in c) shows low phase contrast, consistent with reduced signal due to refractive index matching between tissue and resin. **(e)** Mean *±* s.d. of multiple (n = 25) measurements in this dataset. This dataset contributes an additional data point in Figure 5.

